# A novel fully-automated system for lifelong continuous phenotyping of mouse cognition and behaviour

**DOI:** 10.1101/2022.06.18.496688

**Authors:** Hinze Ho, Nejc Kejzar, Hiroki Sasaguri, Takashi Saito, Takaomi C. Saido, Bart De Strooper, Marius Bauza, Julija Krupic

## Abstract

Comprehensive ethologically-relevant behavioural phenotyping in rodent experiments is essential for deciphering the neural basis of animal cognition. Automated home-cage monitoring systems present a valuable tool to fulfil this need. However, they often involve complex animal training routines, water or food deprivation, and probe a limited range of behaviours. Here, we present a new fully automated AI-driven home-cage system for cognitive and behavioural phenotyping in mice. The system incorporates spontaneous alternation T-maze, novel-object recognition and object-in-place recognition tests combined with monitoring of an animal’s position, water consumption, quiescence and locomotion patterns, all carried out continuously and simultaneously in an unsupervised fashion over long periods of time. Mice learnt the tasks rapidly without any need for water or food restrictions. We applied ethomics approach to show that combined statistical properties of multiple behaviours can be used to discriminate between mice with hippocampal, medial entorhinal and sham lesions and accurately predict genotype of Alzheimer’s disease mouse models on an individual animal level, surpassing the performance of several gold standard cognitive tests. This technology could enable large-scale behavioural screening for genes and neural circuits underlying spatial memory and other cognitive processes.

## Main

Characterising animals’ naturalistic behaviours is paramount to understand how the brain works. Currently, available tests are limited in scope and duration and often lack ethological relevance. In addition, the results commonly show significant variability due to variation in experimental conditions (e.g. food or water restriction, time of an experiment, etc.), individual animal differences and potential subjective bias of the Experimenter^1^. To address these limitations, several automated platforms have been introduced^2–4^. While progress in passive monitoring of motor behaviour (e.g. an animal’s position, speed, posture) has been significant due to advances in software algorithms^3,5,6^ and hardware equipment, extensive ethologically-relevant cognitive phenotyping still presents a major challenge. Namely, all current commercial systems require elaborate pre-training involving food^2,4,7^ or water^2,8,9^ restriction. These requirements not only induce biologically unnatural conditions but can also limit the duration for which the testing may be carried out. Lastly, the current systems are limited in the range of cognitive performances they are designed to test^1^.

Here, we describe a novel home-cage monitoring system called the smart-Kage for fully automated AI-driven comprehensive cognitive and behavioural phenotyping in individually-housed mice, compatible with long-term experiments. To demonstrate the usefulness of this system for basic and translational research, we characterised a small group of mice with hippocampal and medial entorhinal lesions, known to exhibit substantial impairments on spatial memory tasks^10,11^, as well as a widely used *App*^*NL-G-F*^ Alzheimer’s disease mouse model^12,13^. Cognitive tasks include T-maze-like spontaneous alternation (‘smart T-maze’), novel object recognition (‘smart NOR’) and object-in-place recognition (‘smart OPR’) tasks carried out continuously and simultaneously in the smart-Kage without any interference from the Experimenter. In addition, the system monitors an animal’s position, water consumption, quiescence and locomotion patterns. We show that animals with hippocampal lesions can be clearly separated from animals with medial entorhinal lesions and sham controls on an individual animal basis. Moreover, in tandem with a short (∼7 days) test on a standard forced-choice T-maze task, individual mice from all three groups could be separated with an unprecedented >90% accuracy. Finally, we predicted the identity of individual *App*^*NL-G-F*^ mice with 80% (4/5 mice) accuracy in a blind, unsupervised test, surpassing the performance of the gold standard T-maze, NOR and OPR tests.

## Results

### The smart-Kage system

The smart-Kage consists of three connected compartments (two corridors and an open space compartment) separated by three transparent boundaries (Fig. 1a-b). Each corridor leads to a water spout accessed through a nose-poke port with a pair of infrared sensors to detect the mouse’s drinking attempts. The smart T-maze task requires the mouse to alternate between the left and the right corridors to activate the water release, probing the mouse’s ability to recall the position of its previous choice. The water is supplied for as long as the mouse keeps its nose in the port (short <1 s withdrawals were allowed before shifting the active spout to the opposite side). Inter-trial intervals (ITIs) were measured as the time elapsed between spout visits.

**Fig. 1:**
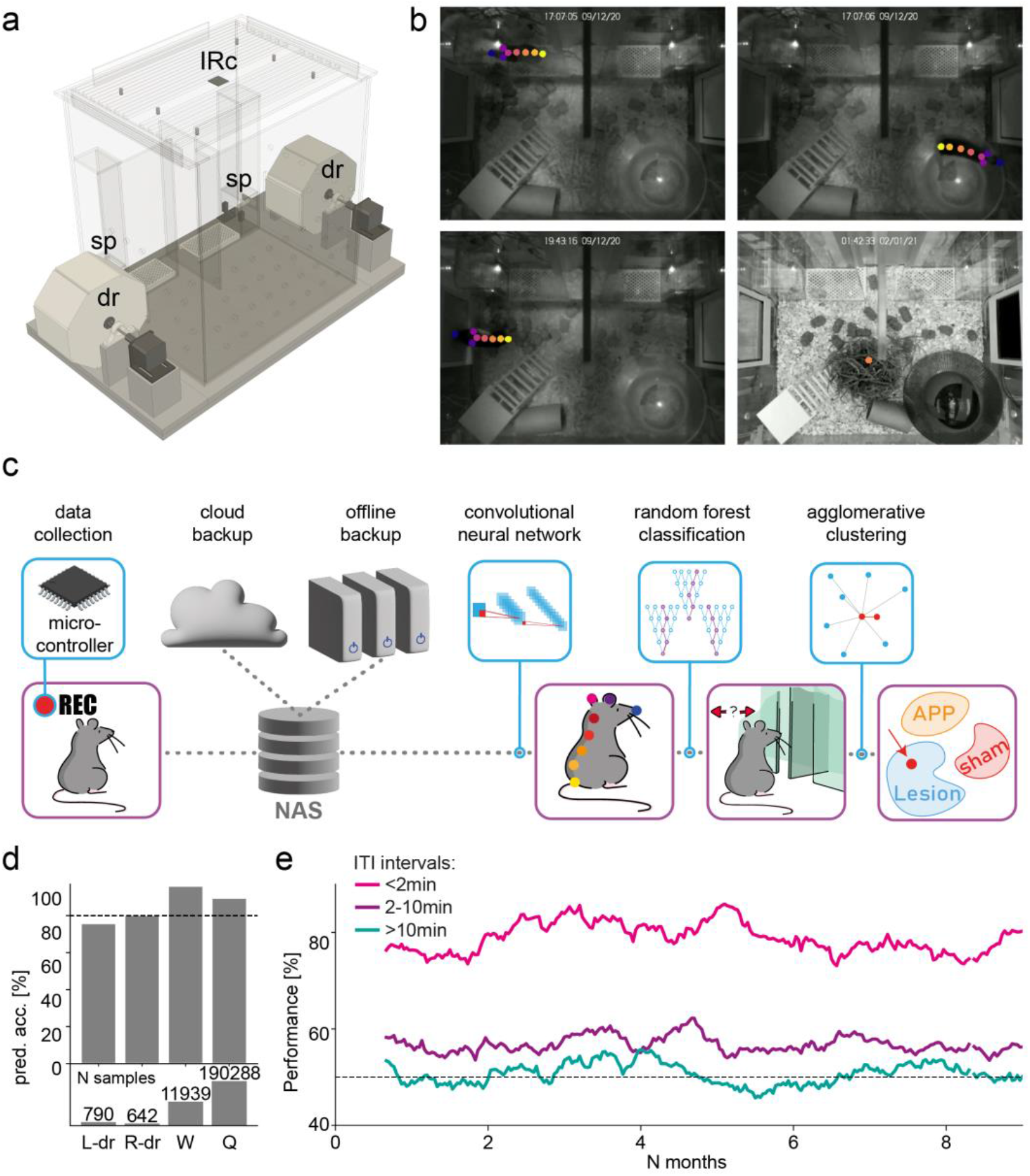
The smart-Kage system. **(a)** Side view of the smart-Kage. IRc: overhead infrared camera; dr: drum; sp: water spout. **(b)** Top views of the smart-Kage interior with example CNN-based video tracking of different mouse behaviours: mouse drinking (top left); mouse running on the wheel (top right); mouse exploring the surface of a side panel (bottom left); mouse in a quiescence state inside the nest (bottom right). Each dot labels a specific mouse body part. **(c)** Automated phenotyping pipeline. Phenotyping begins by collecting top-view videos of the smart-Kage interior through an IR camera. The data is stored on NAS devices and backed up to the cloud and offline external storage. The videos are then analysed using CNN to obtain mouse trajectories and body postures. A random forest classifier is used to assign behavioural labels. In the final stage, a set of behavioural parameters is used to predict the underlying mouse phenotype using agglomerative (hierarchical) clustering. **(d)** The smart-Kage behavioural labelling accuracy (top) is evaluated as the ability of the classifier to avoid false and find true positives (f1 score). The number of ground truth frames used to calculate the phenotyping performance is shown in the bottom; drum exploration (L-dr, R-dr) is a much sparser behaviour compared to running on the wheel (W) or quiescence states (Q). Each T-maze trial prediction was immediately checked against the ground-truth activation of infrared sensors in the spout nose ports, resulting in 100% accuracy in this category (not shown). **(e)** An example of the performance on the smart T-maze at different ITI intervals collected from one mouse continuously tested over >8 months.

The open-space compartment incorporates a continuous novel object recognition task presented as two rectangular surface panels (6.4 × 9.2cm) positioned symmetrically on each side wall (Fig. 1a). The surface panels were made of distinctly different textures and colours, and a mouse could directly explore them by touch, smell and vision. The surface panels are attached to two octagonal drums flanking the sides of the smart-Kage. Different surface panels were presented via the rotation of the drums. The rotation occurred only when the animal was engaged with the water spouts, so it could not directly observe the change. The drums were set to rotate once every two days in around the middle of the dark cycle. The smart NOR task is quantified by measuring the exploration time associated with the change of the surface panel. The changes can happen on either wall individually (left or right NOR) or on both walls simultaneously (double NOR). The smart OPR task consists of keeping the current patterns but ‘swapping’ their locations between the left and right walls. We also implement hybrid changes with one of the walls assuming the pattern identical to the opposite wall while the latter changes to a completely new unseen pattern or remains unchanged (Supplementary Table 1).

Mouse activity within the smart-Kage was continuously recorded by an overhead infra-red (IR) camera (Fig. 1a). The mouse’s precise position was determined by employing a deep convolutional neural network (CNN)^6^ with spatial and temporal resolutions of 0.15 mm and 0.5 s, respectively (Fig. 1a-c; Extended Data Fig. 1; Methods). Mouse behaviours were grouped into four distinct categories of interest (Fig. 1b, Supplementary Videos 1-4): 1) T-maze choices; 2) drum panel exploration; 3) running on the wheel and 4) quiescence state. The behaviours of interest were inferred from the collected set of mouse trajectories and body postures using a random forest classifier^14^, which achieved a >80% prediction accuracy when compared against manually-annotated ground-truth frames (Fig. 1d). All behavioural and cognitive phenotyping tests were run automatically, continuously and in parallel without any interference from the Experimenter. Mice quickly learned the tasks without any need for water or food restriction. Importantly, the system performance was stable over time and was well-suited for long-term studies (Fig. 1e; >8 months, limited only by the duration of the experiment).

### System application for phenotyping different mouse groups

To demonstrate the usefulness of the smart-Kage for cognitive and behavioural phenotyping, we first tested three groups of animals with Experimenter blinded to phenotype: mice with ibotenic-acid-induced lesions in 1) the hippocampus (HP mice); 2) medial entorhinal cortex (mEC mice); and 3) a control group (control mice) that received sham surgical procedures in the hippocampus, medial entorhinal cortex, or medial prefrontal cortex (Extended Data Fig. 2; Supplementary Table 2). All sham groups were combined into a single control group for further analysis since there was no detectable behavioural or cognitive difference amongst them. All animals were randomly assigned to each group prior to commencing the study. The mice were run in two different batches (Supplementary Fig. 1). They were tested for ∼1 month in the smart-Kages before lesioning, followed by additional two months of post-surgery testing.

To benchmark the performance of the smart-Kage, the mice also underwent a battery of gold-standard spatial memory tests (T-maze forced alternation, NOR and OPR tasks)^10,15–18^ before (first batch only) and after (both batches) they were tested in the smart-Kages (Supplementary Fig. 1).

Finally, to demonstrate the usefulness of the smart-Kage for translational research, we also blindly tested a small number of *App*^*NL-G-F*^ mice (Supplementary Table 3), which were previously reported to exhibit mild cognitive deficits on spatial memory tasks^13,14^.

### Smart T-maze task

Since access to the waterspouts was unrestricted, animals were free to choose their own inter-trial intervals (ITIs). For post-analysis purposes we have blocked these ITIs into three groups: <2 min, 2-10 min, and >10 min reflecting short, mid-range and long term working memory, respectively. All pre-lesioned mice rapidly learned the smart T-maze task, performing above chance levels after one day at <2 min inter-trial intervals and after ∼2.5 days at 2-10 min ITIs (Fig. 2a). The performance dropped rapidly with longer ITIs, reaching the chance level at ∼10 min ITI (Fig. 2b), consistent with the working memory time span measured on the standard T-maze task^19^. The spatial working memory was significantly impaired in mice with hippocampal lesions compared to their pre-lesion performance and sham controls (Fig. 2c-d). Specifically, we observed a significant drop in maximum performance (Fig. 2d: 81.8±2.6% pre-lesion vs. 58.3±2.6% post-lesion, U=25, P=0.024, Mann-Whitney U-test) and non-significant but noticeable reduction in the working memory time span (Fig. 2d: 10.2±1.2 min pre-lesion vs. 5.6±1.7 min post-lesion, U=22, P=0.167, Mann-Whitney U-test). On the contrary, there was no significant difference in maximal performance in mice with medial entorhinal lesions (Fig. 2e: 80.3±1.9% pre-lesion vs. 75.1±1.9% post-lesion, U=8, P=1.000, Mann-Whitney U-test). Notably, in HP mice, the performance started to improve after ∼1.5 months post-lesion (Fig. 2c). This performance compensation time scale is comparable to the observations in other well-known hippocampal-dependent tests such as the Morris water maze (∼43 days)^20^.

**Fig. 2:**
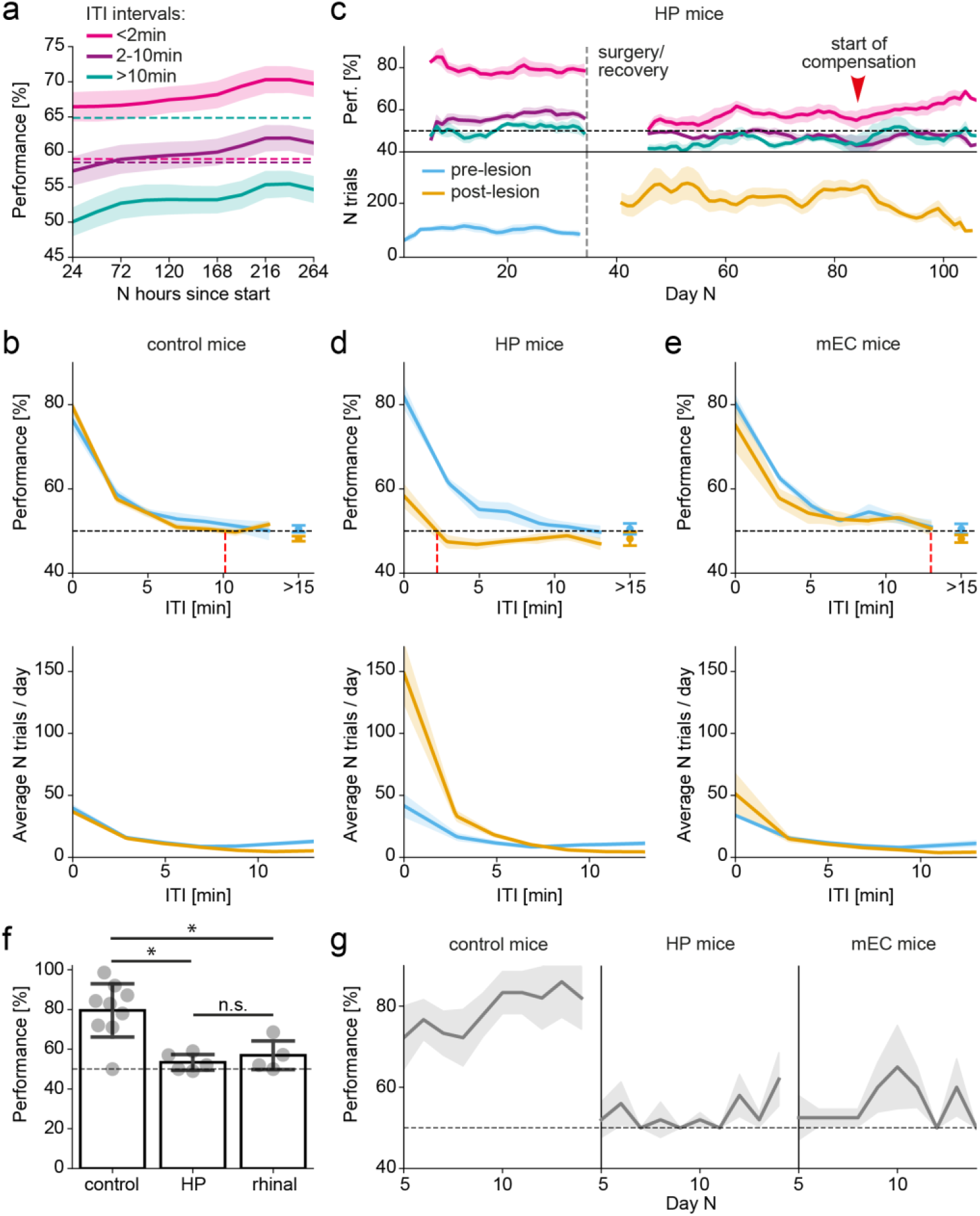
Smart T-maze spatial alternation task. **(a)** The smart T-maze task is rapidly learned at shorter ITI periods (<10 min); short ITI (<2 min): magenta; medium ITI (2-10 min): purple. The performance on longer ITI periods (>10 min) is below chance levels (green). Chance levels (dashed lines) denote the upper bound of the 95% confidence interval of a chance level calculated for the finite number of samples within each ITI domain. **(b)** The average distribution of performance (top) and daily frequency of spout visits (bottom) pre- (blue) and post- (orange) surgery of control mice. The red dashed line indicates the memory time span. **(c)** Running average performance at different ITI intervals (top) and frequency of spout visits (bottom) in HP mice before and after the hippocampal lesions (vertical dashed line). The performance for all ITI domains significantly dropped after the lesion, accompanied by a significant increase in spout visits. The performance started to improve after ∼1.5 months post-lesion (red arrowhead). Horizontal dashed line indicates chance level performance. **(d-e)** The average distribution of performance (top) and daily frequency of spout visits (bottom) in HP (d) and mEC (e) mice, respectively. The red dashed line indicates the memory time span. **(f)** The performance of the same mice on the standard T-maze task. The performances between control and test groups are significantly different (control vs. HP: U=56.5, P=0.018; control vs. mEC: U=44.5, P=0.023; Mann-Whitney U-test), whereas there is no significant difference between HP and mEC mice (U=7.0, P=0.537, Mann-Whitney U-test). **(g)** Daily average performance on a standard T-maze task in control (left), HP (middle) and mEC (right) mice. Shaded regions: standard error of the mean (SEM).

We also found that impairments on the smart T-maze task in HP mice were accompanied by a significant increase in spout visits (Fig. 2d: 41.7±8.9 visits/day pre-lesion vs. 148.4±8.9 visits/day post-lesion, U=1, P=0.048, Mann-Whitney U-test). The results are consistent with previous findings demonstrating larger water consumption in animals with hippocampal lesions^21^. This increase was also accompanied by a higher nose-poking frequency (the mouse repeatedly inserting its head into the spout chamber during the same T-maze trial), suggesting that the mice may have exhibited a compulsive behavioural phenotype (Extended Data Fig. 3: 50.7±8.4 pokes/day pre-lesion vs. 246.6±69.1 pokes/day post-lesion, U=1, P=0.048, Mann-Whitney U-test). No such phenotypes were observed in mice with entorhinal lesions (Fig. 2e, Extended Data Fig. 3: 33.9±0.9 visits/day pre-lesion vs. 51.2±0.9 visits/day post-lesion, U=8, P=1.000, Mann-Whitney U-test; 55.5±3.4 pokes/day pre-lesion vs. 68.2±29.2 pokes/day post-lesion, U=12, P=0.343, Mann-Whitney U-test).

Importantly, unlike the difference in performance between the HP and mEC groups observed on the smart T-maze task, mice with both hippocampal and entorhinal damage showed dramatic impairments on a standard forced-choice alternation T-maze task (Fig. 2f-g). Our finding indicates that although both standard and smart T-maze tasks are hippocampal-dependent, they have remarkably different sensitivity to mEC lesions. Currently, the underlying cause of this difference remains unknown. However, this offers an unprecedented opportunity to use both tests in tandem to distinguish between the hippocampal, medial entorhinal and control animals on an individual animal basis with 94% accuracy.

### Smart NOR and OPR tasks

Animals naturally tend to direct increased exploratory activity towards new stimuli^22^. This observation serves as a basis for some of the most widely used memory tasks in rodent experimentation: novel object recognition (NOR) and object-in-place recognition (OPR) memory tasks^16,17,23^. It has been suggested that the performance on standard NOR and OPR tasks may be affected in animals with parahippocampal and hippocampal lesions, respectively^11^. However, the findings are often heavily influenced by the exact experimental procedures^23^. To overcome this difficulty, we implemented analogous tasks in the smart-Kage.

Our results show that all animal groups increased their exploration time in response to the change of a surface panel (Fig. 3a-b). Similar to observations in standard tests, the increase in exploration was a natural behaviour which did not require any pre-training and occurred from the first encounter (Extended Data Fig. 4). The time to notice the change was comparable in all mice groups (Fig. 3b; 1.2±0.1min vs 1.6±0.2min vs 1.4±0.2min in control, HP and mEC groups, respectively; F=2.58, P=0.109, one-way ANOVA). On the other hand, the average exploration time in both smart NOR and OPR tasks tended to be longer in mice with hippocampal lesions (Fig. 3d: 5.2±1.1 min/day pre-lesion vs. 11.1±1.8 min/day post-lesion, U=2, P=0.095, Mann-Whitney U-test) compared to mEC and control mice (Fig. 3c,e: control mice: 4.6±0.4 min/day pre-lesion vs. 4.6±0.4 min/day post-lesion, U=41, P=1.000, Mann-Whitney U-test; mEC mice: 6.4±0.7 min/day pre-lesion vs. 6.2±1.1 min/day post-lesion, U=9, P=1.0000, Mann-Whitney U-test), which remained similar to pre-surgical levels. This difference may reflect the increased compulsivity in HP animals, similar to the increase in drinking behaviour observed on the smart T-maze task. Importantly, we found no significant difference between all groups when tested on the standard NOR and OPR tasks (Extended Data Fig. 5; Methods).

**Fig. 3:**
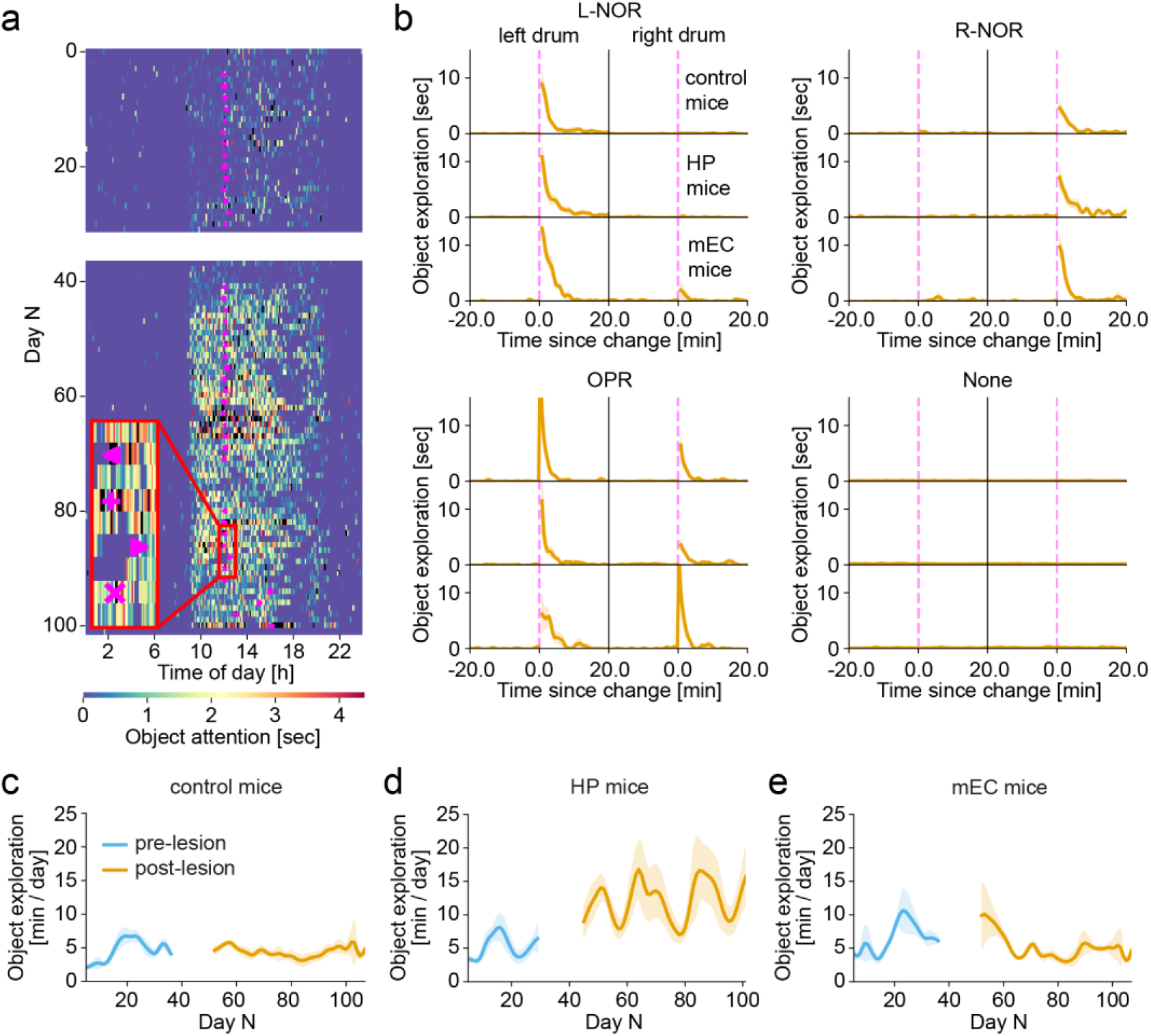
Smart novel object and object-in-place recognition tasks. **(a)** An example of wall panel exploration patterns of a single HP mouse tested for ∼3 months. The time of lesion surgery and the following week of recovery are indicated by the white gap between days 30 and 40. Purple shapes indicate the time of side-panel change (left-facing triangle: left drum NOR; plus: left drum OPR + right drum NOR; right-facing triangle: right drum NOR; X: left drum NOR + right drum OPR). (**b**) Animal response to L-NOR (top left), R-NOR (top right) and OPR (bottom left) tasks 20 min before and after the change of a side panel, indicated with the purple dashed line. Top, middle and bottom rows of each plot correspond to control, HP and mEC mice, respectively. Left and right columns of each plot correspond to the exploration time of the left and the right side panel, respectively. Bottom left and right plots show animal exploration time when both or none of the side panels changed. The exploration at no change was measured at 12 pm for direct comparison. **(c-e)** Average daily side panel exploration time for control (c), HP (d) and mEC (e) mice, respectively. Pre-lesion period: blue; post-lesion period: orange; shaded regions: standard error of the mean (SEM).

### Locomotion and quiescent behaviors

In addition to measuring animals’ performance on cognitive memory tasks, we also assessed their locomotion and quiescence patterns. Voluntary activity on the running wheel has been previously associated with increased hippocampal neurogenesis and improvements in animal performance on spatial memory tasks^24,25^. However, it is unknown how mice with hippocampal and medial entorhinal damage use a running wheel. Our results show that mice with hippocampal damage showed ∼67% reduction in time spent running on the wheel (Fig. 4a-b: 185.5±40.5 min/day pre-lesion vs. 60.9±19.5 min/day post-lesion, U=21, P=0.286, Mann-Whitney U-test) in contrast to a significant ∼78% increase in their overall movement in the smart-Kage (Fig. 4b: 385.5±19.1 min/day pre-lesion vs. 685.3±31.5 min/day post-lesion, U=0, P=0.024, Mann-Whitney U-test). Interestingly, the increase in movement was accompanied with instances of stereotypical behaviours, such as running in a circular trajectory which was not observed in control animals (Supplementary Video 5). No decrease in running on the wheel was observed in mice with medial entorhinal lesions (Fig. 4b: 127.7±16.6 min/day pre-lesion vs. 141.1±9.7 min/day, U=7, P=0.89, Mann-Whitney U-test) or control group (Fig. 4b: 133.4±20.9 min/day pre-lesion vs. 133.1±12.5 min/day post-lesion, U=46, P=0.886, Mann-Whitney U-test). They also did not exhibit any noticeable change in the overall movement in the smart-Kage post-surgery (Fig. 4b: mEC mice: 445.1±19.0 min/day pre-lesion vs. 498.6±25.4 min/day post-lesion, U=5, P=0.486, Mann-Whitney U-test; control mice: 420.2±24.1 min/day pre-lesion vs. 454.6±31.6 min/day post-lesion, U=30, P=0.486, Mann-Whitney U-test).

**Fig. 4:**
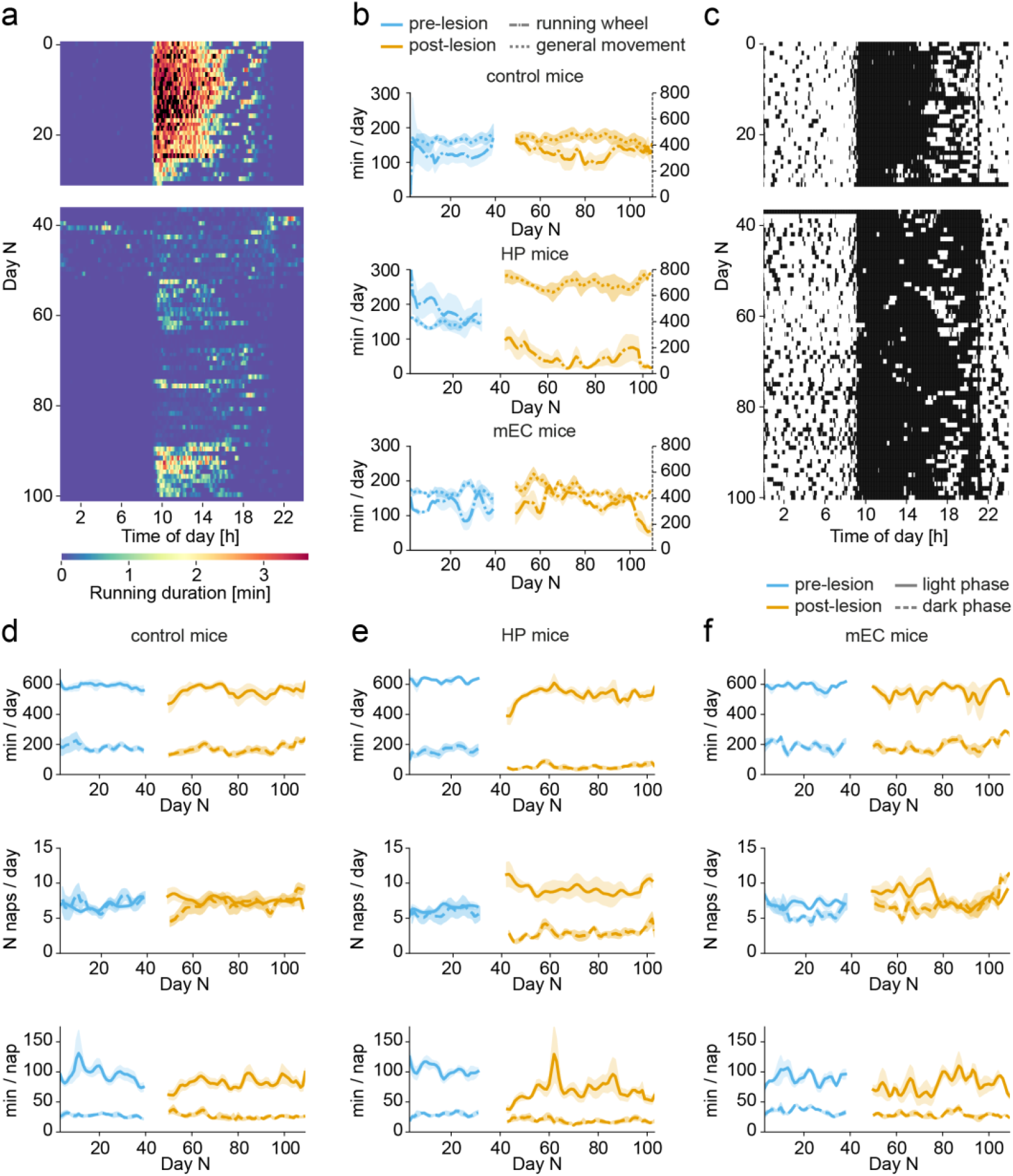
Monitoring locomotion and quiescence states in the smart-Kage. **(a)** An example ethogram showing daily wheel-running activity of a mouse with a hippocampal lesion. The time of lesion surgery and the following week of recovery are indicated by the white gap between days 30 and 40. **(b)** Average general locomotion (dotted) and wheel-running (dash-dotted) behaviours in control (top), HP (middle) and mEC (bottom) mice, respectively. **(c)** An example ethogram showing quiescence states of the same mouse as in (a). White and black regions indicate quiescence and mobile intervals, respectively. The time of lesion surgery and the following week of recovery are marked as the white gap between days 30 and 40. **(d-f)** The total daily average time spent in quiescence (top), daily average number of quiescence states (middle) and their average duration (bottom) in control (d), HP (e) and mEC (f) mice. Solid and dashed lines correspond to light- and dark-phases, respectively. Pre-lesion period: blue; post-lesion period: orange; shaded regions: standard error of the mean (SEM).

Next, we assessed the animal’s quiescent states, defined as periods when the animal was completely and continuously motionless for at least 5 min (Methods) and as such, they were used as a proxy for estimating sleep patterns. We found that control animals spent most of their 12-hour light cycle immobile or sleeping (Fig. 4c-d: 596.1±15.3 min/day pre-lesion vs. 561.3±17.2 min/day post-lesion, U=60, P=0.140, Mann-Whitney U-test) with occasional brief activity periods usually related to water or food consumption. Moreover, control animals often developed regular quiescence patterns during the dark cycle (Fig. 4c-d: 179.5±15.3 min/day pre-lesion vs. 161.9±13.9 min/day post-lesion, U=51, P=0.566; 6.8±0.6 states/day pre-lesion vs. 6.5±0.8 states/day post-lesion, U=47, P=0.596; 28.4±1.2 min/state pre-lesion vs. 29.3±3.0 min/ state post-lesion, U=37, P=0.791, Mann-Whitney U-test). On the other hand, mice with hippocampal lesions showed significantly disrupted and less regular quiescence patterns (Fig. 4c, e) characterised by a significant decrease in average immobility time (Fig. 4e: light phase: 625.4±7.1 min/day pre-lesion vs. 524.1±11.1 min/day post-lesion, U=25, P=0.024; dark phase: 155.4±21.0 min/day pre-lesion vs. 52.6±7.2 min/day post-lesion, U=25, P=0.024 Mann-Whitney U-test); and significantly increased frequency of short quiescence states during the light phase (Fig. 4c, e: 6.4±0.2 states/day pre-lesion vs. 9.5±0.9 states/day post-lesion, U=0, P=0.024, Mann-Whitney U-test). Generally, the quiescence became less regular

The mice with medial entorhinal lesions showed smaller and more transient changes in immobility patterns, characterised mostly by the non-significant increase in the number of shorter quiescence states during the light cycle (Fig. 4f: 7.0±0.4 states/day pre-lesion vs. 9.1±1.0 states/day post-lesion, U=4, P=0.343; 91.2±4.3 min/state pre-lesion vs. 69.9±13.2 min/state post-lesion, U=12, P=0.343, Mann-Whitney U-test), which returned to normal levels in about a month.

### Automated classification of individual animals

Can we use the smart-Kage to classify different mouse strains on an individual animal basis? To address this question, we blindly tested four different mouse groups: *App*^*NL-G-F*^ Alzheimer’s disease mouse model group as well as HP, mEC and control groups (which included both sham lesion controls and *App*^*NL-G-F*^ negative mice). Multiple cognitive and behavioural measurements (Supplementary Table 3) were grouped together using an agglomerative (hierarchical) clustering algorithm to predict the underlying mouse phenotype (Methods). Specifically, we ran >10,000 clustering simulations to optimise the clustering hyper-parameters and determine the best combinations of cognitive and behavioural measurements as well as the testing periods. The clustering parameters were chosen so that a maximum number of pre-lesioned animals, presumably with a highly similar behavioural and cognitive phenotype, were assigned to a single ‘control’ cluster. Out of the resultant set of clustering parameters, the final optimal clustering was chosen after unblinding to maximise the separation between all different mouse groups (Fig. 5).

**Figure 5:**
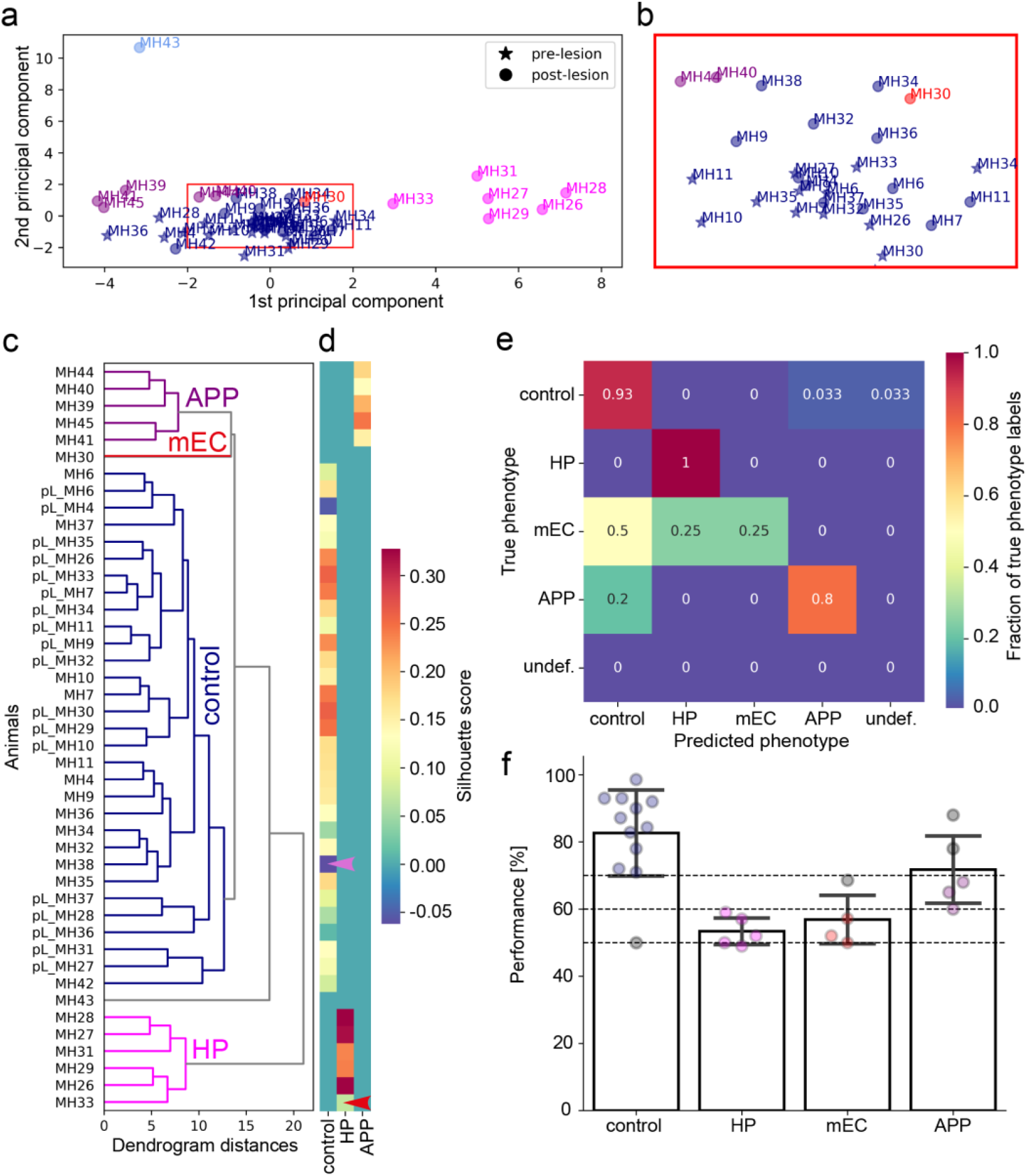
Behavioural clustering on individual animal basis. **(a)** Cluster assignments in the original 32-dimensional feature space visualised with 2D PCA. Different colors correspond to different clusters. The dot labels indicate animal identities. Pre- and post-lesion feature vectors are marked with stars and circles, respectively. **(b)** An inset region, marked in (a). **(c)** Hierarchical clustering dendrogram. Colours correspond to clustering colours in (a). **(d)** Clustering silhouette plot. Warmer colours indicate good separation from neighbouring clusters, whereas colder colours indicate potential outliers in respective clusters. An mEC outlier in the HP cluster and APP outlier in the control cluster are marked with red and purple arrowheads, respectively. **(e)** Confusion matrix comparing actual and predicted animal identities detected by smart-Kage. The values indicate the percentage of animals assigned to each class. **(f)** Animal clustering based on the standard T-maze task. The dashed lines demarcate distinct clusters: controls (blue) at >70% performance, HP and mEC mice at 50-60% performance (pink and red), *App*^*NL-G-F*^ mice at 60-70% performance (purple). Grey circles mark misidentified mice.

Using this approach, we were able to correctly classify 100% (27/27) of sham control mice, which were clustered together with the pre-lesioned mice (i.e. a control cluster), 100% (5/5) of mice with hippocampal lesions and 25% (1/4) of mice with medial entorhinal lesions (Fig. 5a-e). Interestingly, one of the mEC animals was clustered together with the HP group but stood out as an outlier within this group (Fig. 5d, red arrowhead). The two remaining mice with medial entorhinal lesions were misclassified as control. After unblinding of the histology results, we found that the correctly identified mEC mouse had the largest volume of the medial entorhinal cortex removed, followed by the mouse within the HP cluster (Supplementary Table 2). The mEC mice grouped with the control animals had much more limited mEC lesions.

Importantly, we also successfully identified 80% (4/5) of *App*^*NL-G-F*^ mice. The only misassigned *App*^*NL-G-F*^ mouse was classified as a control animal; although it was a strong outlier within the control group (Fig. 5d, purple arrowhead). Of note, *App*^*NL-G-F*^ control animals were classified outside the general control cluster likely due to their different genotype and/or age compared to pre-lesioned animals and sham controls (Methods). Notably, the classification accuracy achieved by the smart-Kage surpassed the standard T-maze, NOR and OPR tasks (Fig. 5f; 60% (3/5) *App*^*NL-G-F*^ mice). In the case of the standard tests, the grouping was based on the performance on the T-maze alone, as results from NOR and OPR tasks did not serve as good predictors (Extended Data Fig. 6; Methods). Adding more tests to the battery, such as open-field exploration could potentially improve the accuracy of standard tests. However, each new test would require separate pre-training and currently, no set of standard tests is able to separate HP and mEC animals.

## Discussion

Here we described a novel home-cage monitoring system (the smart-Kage) which incorporates T-maze, NOR and OPR tests, enabling their autonomous, simultaneous and continuous execution. In the current study, the phenotyping was focused on cognitive domains especially important for research of neurological disorders, but it can also be used to study decision making, compulsivity and many other cognitive trends. We showed that the delay in water consumption encountered after choosing an incorrect water spout was a sufficiently negative reinforcer to trigger rapid learning on the smart T-maze task eliminating any need for food or water restriction and thus contributing to the ethological relevance of our system. In addition, smart-Kage also simultaneously characterises a range of other non-cognitive behaviours such as locomotion and quiescence states, which in the majority of cases likely serve as a proxy for an animal’s sleeping patterns.

In the proof-of-principle experiments using small samples of mice with hippocampal, medial entorhinal and sham lesions as well as *App*^*NL-G-F*^ Alzheimer’s disease mice and their controls, we demonstrated the effectiveness of the smart-Kage by focusing on hippocampal-parahippocampal-dependent spatial working memory, novel object and object-in-place recognition behaviours. In a blind test, applying ethomics approach we showed that using the smart-Kage, we could identify different groups of animals on an individual animal basis with high accuracy without making any assumptions about specific unambiguous group phenotypes. Instead, we relied solely on combining multiple behavioural measures simultaneously recorded in the smart-Kage, which on their own often showed only non-significant behavioural and cognitive trends (at least in the small animal samples that we employed). Unlike previously reported approaches which used the ‘leave-one-animal-out procedure’^7^ (i.e. the identities of all but one animal were provided for cluster assignment), our clustering algorithm was trained only on the pre-lesioned mice representing a known control group. All other clusters, which included both lesion and sham post-lesion groups and mice with Alzheimer’s disease-associated genetic modifications, were produced automatically in a completely unsupervised way; and the best solution was chosen based on the optimal group assignment for all groups combined. We showed that our approach yielded superior results compared to behavioural phenotyping using the three most prominent analogous standard memory tests.

The system’s autonomy and ethological setup eliminate human bias, potentially presenting us with the first real solution to the reproducibility crisis in mouse behaviour research. The widespread adoption of such (or similar) systems across different laboratories would result in the global pooling of directly comparable data and supercharge the future of cognitive and behavioural research.

## Methods

### Subjects

This work was conducted in accordance with the UK Animals (Scientific Procedures) Act (1986). Four groups of mice were used in the study: C57BL/6J mice with lesions to 1) the hippocampus, 2) medial entorhinal cortex or 3) sham controls, with saline injections in the hippocampus, medial entorhinal cortex or medial prefrontal cortex. The fourth group comprised *App*^*NL-G-F*13,14^ mice (Supplementary Table 2).

19 male C57BL/6J mice sourced from Charles River underwent lesion procedures. The experiments were carried out in two Batches ∼3 months apart. The mice were 10-16 weeks old when they were transferred to the smart-Kages. They were individually housed in the smart-Kages for ∼30 days prior- and ∼60 days post-surgery. The mice weighed 25-30 g at the time of the surgery. Water and food were supplied ad libitum. The first batch was tested on the standard forced alternation T-maze task, object recognition and object-in-place recognition tasks before and after testing in the smart-Kages. The second batch underwent the same standard tests only after their testing in the smart-Kages was completed. The mice were individually housed in clear plastic cages (16 cm × 27 cm × 18 cm, W × L × H) when they were tested on standard tasks (Supplementary Fig. 1). They were maintained on 85% body weight food restriction schedule when tested on the standard forced-choice T-maze alternation task.

Three male *App*^*NL-G-F*^ KI male mice^13,14^ and three matched controls were included in blinded test experiments. They were 22-23 weeks old when first tested in the smart-Kages. We also included additional two *App*^*NL-G-F*^ positive males aged 39 weeks whose identity was known. All 8 mice were tested on the standard tests before and after they were tested in the smart-Kages.

All mice were kept on a 12:12 h light:dark cycle (with lights on at 9:00 am and off at 9:00 pm) at a controlled temperature (21–23 °C) and humidity (50–60 %).

### Surgery

Mice were anaesthetised with 1–3% isoflurane in O_2_, and 0.05mg/10g body weight Metacam and 0.05mg/10g Baytril were administered to facilitate recovery. Chemical lesions were induced by injection of 10µg/µl ibotenic acid dissolved in pH7.4 PBS into selected brain regions using Nanofil syringe controlled by the micropump. To induce hippocampal lesions, we used the same coordinates and injected volumes as previously described in^9^. To induce mEC lesions, we aimed to bilaterally inject in the following four coordinates. mEC1 (150 nl): AP: 0.4 mm anterior to sinus; ML: 3.4 mm from the midline; DV:2.4 mm; mEC2 (150 nl): AP: 0.4 mm anterior to sinus; ML: 3.4 mm from the midline; DV:1.6 mm; mEC3 (150 nl): AP: 0.4 mm anterior to sinus; ML: 2.8 mm from the midline; DV:3.0 mm; mEC4 (150 nl): AP: 0.4 mm anterior to sinus; ML: 2.8 mm from the midline; DV:2.2 mm. The injection syringe was tilted at 6 degrees anterior-to-posterior angle. Following surgery, mice were individually housed in a conventional cage, and their health conditions were monitored for 6 days before they returned to their corresponding smart-Kage when fully recovered.

### Blinding procedures

The brain regions targeted for lesioning were known only to the researcher who conducted the lesion surgeries. Mice were selected at random and their identities were unknown. Experimenters, who were responsible for conducting behavioural tests, maintenance of smart-Kages, data collection and analysis, were blinded to the lesion identity and groups for the duration of the experiments. The genotypes of three homozygote *App*^*NL-G-F*^ mice and three control littermates were kept hidden from all of the Experimenters until the analysis was completed. The identities of two older homozygote *App*^*NL-G-F*^ mice were known to the Researchers.

During the lesion quantification from the histology, the Experimenter was blind to the animals’ characterization in the smart-Kage or their performances on the standard tests.

### Histology

Following completion of the experiments, the mice were given an overdose of sodium pentobarbital and perfused transcardially with phosphate-buffer saline (PBS), followed by 4% formaldehyde to fixate the brain tissue. The brains were carefully extracted from the skull and stored in 4% paraformaldehyde (PFA) at 4°C. Brains were then imaged using serial two-photon tomography^26^ which sliced and imaged the entire brain every 20 μm coronally with a resolution of 4 μm in x and y using autofluorescence at 800 nm. To estimate mEC lesions, brains were resliced computationally into sagittal sections. Lesion volume was estimated by manually marking the total brain area volume and the lesion volume every 60 μm (for HP lesions) and 50 μm (for mEC lesions).

### Smart-Kage design

The smart-Kages and associated components were designed using computer-aided design software. The smart-Kages were manually assembled from parts made of 3- and 5-mm thick transparent acrylic sheets that were laser-cut into correct dimensions and designs. The final dimensions of smart-Kage were 39 cm x 32 cm x 44 cm (W x L x H). The smart-Kage was fastened to a base and flanked by two drums used for NOR and OPR tasks (see above). An overhead infra-red camera was installed on a removable lid, providing a top view into the Kage interior and was used for continuous video recording at two frames/second. Infra-red LEDs were distributed around the lid to provide illumination. The smart-Kage was fitted with sensors and electronic accessories, all connected to and controlled by a single-board microcontroller. All data generated was automatically transferred to a single-board computer for data sorting and storage (see below). The smart-Kage contained a running wheel and other enrichment items (e.g. a climbing platform and a nesting material). The smart-Kage included three integrated cognitive tasks: the smart spontaneous T-maze task, smart NOR and OPR tasks. Each mouse interacted with the tasks on its own volition and data was continuously gathered using sensors and video camera.

### Experimental procedures

Mice were group-housed and acclimatised in the holding facility for at least one week prior to the start of experiments. Before transferral to the smart-Kages, the first batch of C57BL/6J and *App*^*NL-G-F*^ mice underwent a series of standard behavioural tasks, comprising forced-choice alternation T-maze task, novel object recognition (NOR) and object-in-place recognition (OPR) tests. All mice participated in the same set of tests following the smart-Kage experiment. The standard testing protocols were based on published versions in the literature^17,18^ and briefly described below.

### Testing in the smart-Kages

Mice were kept single housed in the smart-Kages with free access to food and water. During the habituation stage (5-7 days), mice received water from both water spouts. After the habituation period, a smart spontaneous alternation T-maze task commenced when only one of the water spouts (an active spout) provided water at any given time. The location of the active water spout alternated every time a mouse received the water (i.e. accessed an active spout). The smart NOR and OPR tests were implemented by rotating the side drums to present one of the eight side patterns (0.5 cm x 6.4 cm x 9.2 cm) accessible for a mouse to explore. The drum rotation was programmed to occur every two days between 12 am – 4 pm and occurred only when a mouse was drinking water in one of the corridors to ensure that it could not observe the change. The patterns were presented according to a schedule designed to test both spatial and non-spatial ‘object’ recognition abilities of the mouse. The drum-change sequence consisted of the following combinations: (1) left-drum only NOR, (2) right-drum only NOR, (3) both drums NOR, (4) left-drum only OPR, (5) right-drum only OPR, (6) both drums OPR, (7) left-drum NOR/right-drum OPR, and (8) right-drum NOR/left-drum OPR.

Mice underwent lesion surgery after 4-6 weeks of residing in the smart-Kages and were kept in conventional cages for 6 days of post-operative care. Following full recovery from surgical procedures, mice were transferred back to the same smart-Kages and were kept for further 8-12 weeks. The smart-Kages were cleaned every two weeks. Mice were tested in standard behavioural tests following the smart-Kage experiment, as described below.

### Standard forced-choice alternation T-maze task

The test was conducted using a T-shaped enclosure consisting of a start arm adjoining two perpendicular goal arms. Mice were food-restricted for at least 12 h before each experiment day and were kept at approximately 90% of initial body weight. Soya milk was used as a reward and was located in a food well at the end of goal arms. One day prior to the testing session, mice were habituated to the apparatus by allowing them to freely explore the enclosure. The habituation consisted of four 3-minutes periods of exploration, interleaved by a 10-minute interval. During the habituation, mice could drink soya milk *ad libitum* from the food well at both goal arms.

Each daily session was composed of 10 trials, and each trial consisted of a sample run followed by a test run. Each pair of the sample and test runs were separated by about 12-15 min. During the test run, the animal was kept in the start arm for 10 seconds before it was allowed to explore the goal arms. If the goal arms were not visited within 90 s, the animal was removed and the trial was terminated.

### Standard NOR and OPR tests

We closely followed the protocol described in^17^. The tests were performed in a 0.5 × 0.5m^2^ square enclosure. Objects were placed 15 cm from the walls. Object exploration was defined as instances when the mouse looked or sniffed at the object in proximity (<2 cm) or when there was direct contact with a snout or paws. Climbing or chewing was not counted as object exploration. One day before the tests began, the mice were allowed to freely explore the enclosure without any objects present. The habituation consisted of two 10-minutes periods of exploration, with a 3-hour interval in between. The tests comprised a familiarisation session followed by a test session with a two-hour inter-trial delay. For NOR, two objects were included in each session, and for OPR, two pairs of objects were used. The familiarisation session was run for 5 minutes unless a mouse explored an object for over 40 s, at which point the mouse was removed and the trial terminated. In NOR, one of the two objects was replaced with a new one between sessions. The other object was also replaced with an identical object to ensure that familiarity was not based on an animal marking the familiar object. In the OPR task, all of the objects were replaced with identical objects during the test trial, and the positions of one pair were swapped between sessions. An overhead camera was used to capture videos of mouse activity for post-hoc visual inspection and analysis.

### Smart-Kage data collection

Parameters and functions of smart-Kage sensors and motors were configured using a custom-written script. An overhead IR camera was used to capture videos of mouse activity. Additionally, every approach towards the water spouts (both correct and incorrect choices) was relayed by beam-break sensors to the controller. Time was synced directly from the internet. All data generated was instantaneously transmitted to a computer for storage, logging and sorting.

### Mouse video tracking

Animal location was tracked from recorded videos using a ResNet-101 deep convolutional neural network (CNN). The starting architecture (pre-trained on ImageNet) was retrained for mouse tracking within the smart-Kage using transfer learning implemented in DeepLabCut^6^ (DLC) software and a training dataset of video frames manually labelled with 8 mouse body parts (snout, left and right ears, neck, 3 points along the mouse’s spine and tail base; Fig. 1b). The retraining was done in 9 consecutive cycles with the ADAM optimiser, batch processing (batch size 8) and *imgaug* image augmentor^6,27^. In each consecutive training cycle, the training dataset was manually expanded with frames on which the resulting network from the previous cycle performed poorly. The cycles were repeated until the test dataset error (MAE between predicted and ground truth mouse body-part labels) plateaued in cycle 9 at 1.41 px (0.15 mm). A total of 800 manually labelled frames were used.

### Behavioural labelling

Behavioural labels (e.g. ‘exploring NOR’, ‘T-maze trial’, ‘quiescence’ etc.) were assigned to mouse trajectories and body postures in three main ways. Smart T-maze trials were assigned whenever a mouse presence in the corridors coincided with beam-breaker detection. Sleeping was assigned to frames with little or no detected motion, cross-validated with frame-subtraction. Finally, pattern exploration and running-wheel exercise were assigned with a random forest classifier. The classifier hyperparameters were tuned with randomised 3-fold cross-validation, and the classifier was subsequently trained on a training dataset of video frames manually labelled with ground-truth behavioural labels. The classifier performance for each behavioural category was estimated on the test dataset, as shown in Fig. 1d. A total of 542530 manually labelled frames were used, which includes training dataset and test dataset (Fig. 1d displays the test dataset).

### Smart-Kage prediction on an individual animal basis

Based on the assigned behavioural labels, various statistics were calculated for all behavioural categories (Figs. 2-4). From these, 32 behavioural measures were gathered into feature vectors, specific to each animal (Supplementary Table 3). The feature vectors were standardised to 0 mean and unit variance to address differences in scale between features. To predict underlying mouse phenotypes, a set of feature vectors from all animals was then clustered with agglomerative clustering. Specifically, >10000 clustering simulations were run to account for hyperparameter optimisation and test which combinations of behavioural measures (and over what testing periods) produced the ‘optimal’ clustering. This was defined as clustering, which assigned the maximum number of pre-lesioned animals (the only known group) into a single ‘control’ cluster and maximised the separation between different mouse strains after unblinding. The optimal clustering (using ward linkage and Euclidean metric) yielded 7 clusters (Fig. 5). Cluster outliers were identified with silhouette analysis – a negative silhouette score indicates an outlier^28^.

### Standard test prediction on an individual animal basis

We used the performance measurements from standard T-maze, NOR, and OPR tasks to group the animals on an individual animal basis to benchmark the prediction of the smart-Kage against analogous standard tests. 42.86% (12/28 animals) and 46.43% (13/28 animals) of animals were discarded in NOR and OPR tests, respectively, at a 30% threshold difference between the exploration of both objects during the familiarisation session. The remaining animals were clustered with simple threshold criteria. Similarly to the smart-Kage clustering above, these thresholds were selected to assign a maximum number of control animals into a single ‘control’ cluster. Animals with T-maze performance below 70%; the absolute value of NOR d2 ratio^29^ below 0.04 and the absolute value of OPR d2 ratio below 0.06 were identified as displaying cognitive decline. In brief, d2 ratio is the difference in exploration time between the novel and familiar object, normalized with regard to their combined exploration time. Hence, a d2 value of 0 indicates equal exploration time between the two objects, whereas values closer to +/-1 indicate a preference for one of the objects.

### Statistics

The effects of different lesions were tested by statistically comparing pre- and post-lesion periods of equal time spans (∼30 days) within each test group (control, HP and mEC mice). The mean value of a given measure was calculated for each animal and all combined means were compared with Mann-Whitney’s U test^30^, except when comparing animal reaction times to changes of drum patterns between the 3 lesion groups. In this case, one-way ANOVA was used instead. All P-values were corrected for multiple comparisons using the Benjamini-Hochberg correction. All data presented is reported as mean ± s.e.m. unless stated otherwise.

## Data availability

All data is available at [add link].

## Code availability

Processing scripts are deposited at [add GitHub link].

## Acknowledgment

We thank Marino Krstulovic and Eszter Arany for helping with smart-Kage maintenance. We also thank J. O’Keefe and K. Duff for their comments on the manuscript. We would also like to thank Sainsbury Wellcome Centre Advanced Microscopy Core for access to their brain histology equipment. The *App*^*NL-G-F*^ mice were generated by the RIKEN BRC through the National BioResource Project of the MEXT/AMED, Japan. In blind testing, three *App*^*NL-G-F*^ and three matched control mice were kindly supplied by F. Wiseman and S. Wells from the UK DRI Animal Models Programme at the Mary Lyon Centre Harwell, supported by the UK Dementia Research Institute (UKRI-1019), which receives its funding from DRI Ltd, funded by the UK Medical Research Council, Alzheimer’s Society and Alzheimer’s Research UK. Two *App*^*NL-G-F*^ mice were supplied by the Paulsen lab at the University of Cambridge. This work was supported by Dementia Research Institute (DRICAMKRUPIC18/19). N.K. is supported by MRC DTP at the University of Cambridge. J.K. is a Wellcome Trust/Royal Society Sir Henry Dale Fellow (206682/Z/17/Z) and is supported by Isaac Newton Trust/Wellcome Trust ISSF/University of Cambridge Joint Research Grant, Kavli Foundation Dream Team project (RG93383), Isaac Newton Trust [17.37(t)], and NVIDIA Corporation.

## Contributions

J.K. and M.B. conceived the study, and developed and built the prototype of the smart-Kage. H.H. carried out smart-Kage construction, functionality optimisation, and data collection with contributions from M.B., J.K. and N.K. J.K. performed the lesion surgeries. N.K. developed the data analysis pipeline with contributions from M.B., J.K. and H.H. M.B. did the histology. H.S., T.S. and T.C.S. generated the *App*^*NL-G-*^ mice. J.K., M.B., N.K. and H.H. wrote the manuscript with contributions from B.D.S. J.K and B.D.S acquired funding.

## Corresponding authors

Correspondence to Marius Bauza or Julija Krupic.

## Extended Data

**Extended Data Fig. 1:**
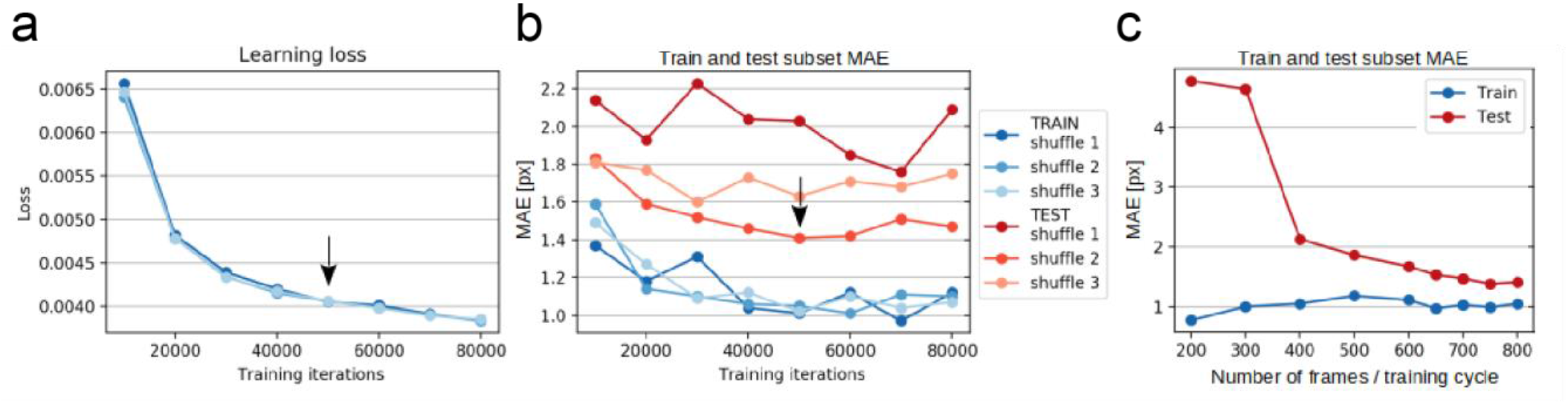
Behavioural clustering on an individual animal basis. **(a)** An example learning-loss plot from a single training cycle with 80k training iterations. The black arrow indicates a plateau in learning loss, at which point the network weights for the subsequent training cycle were taken. **(b)** An example plot from a single training cycle with 80k training iterations showing the network performance evaluation on the train (blue) and test (red) subsets (95% and 5% of the original training dataset, respectively). The performance is measured as the mean absolute Euclidean distance (MAE) between human-annotated and network-predicted body part labels in units of pixels. Note that unlike in root mean square error (RMSE), in MAE, the Euclidean distance is calculated before calculating the mean over frames and body parts (i.e. operations of square root and mean are inverted). The black arrow indicates a minimum in test subset MAE of the shuffle 2, at which point the network weights for the subsequent training cycle were taken. **(c)** Neural network training overview. The network was trained until the test dataset MAE plateaued at 1.41 px (9 cycles, 800 total training frames).

**Extended Data Fig. 2:**
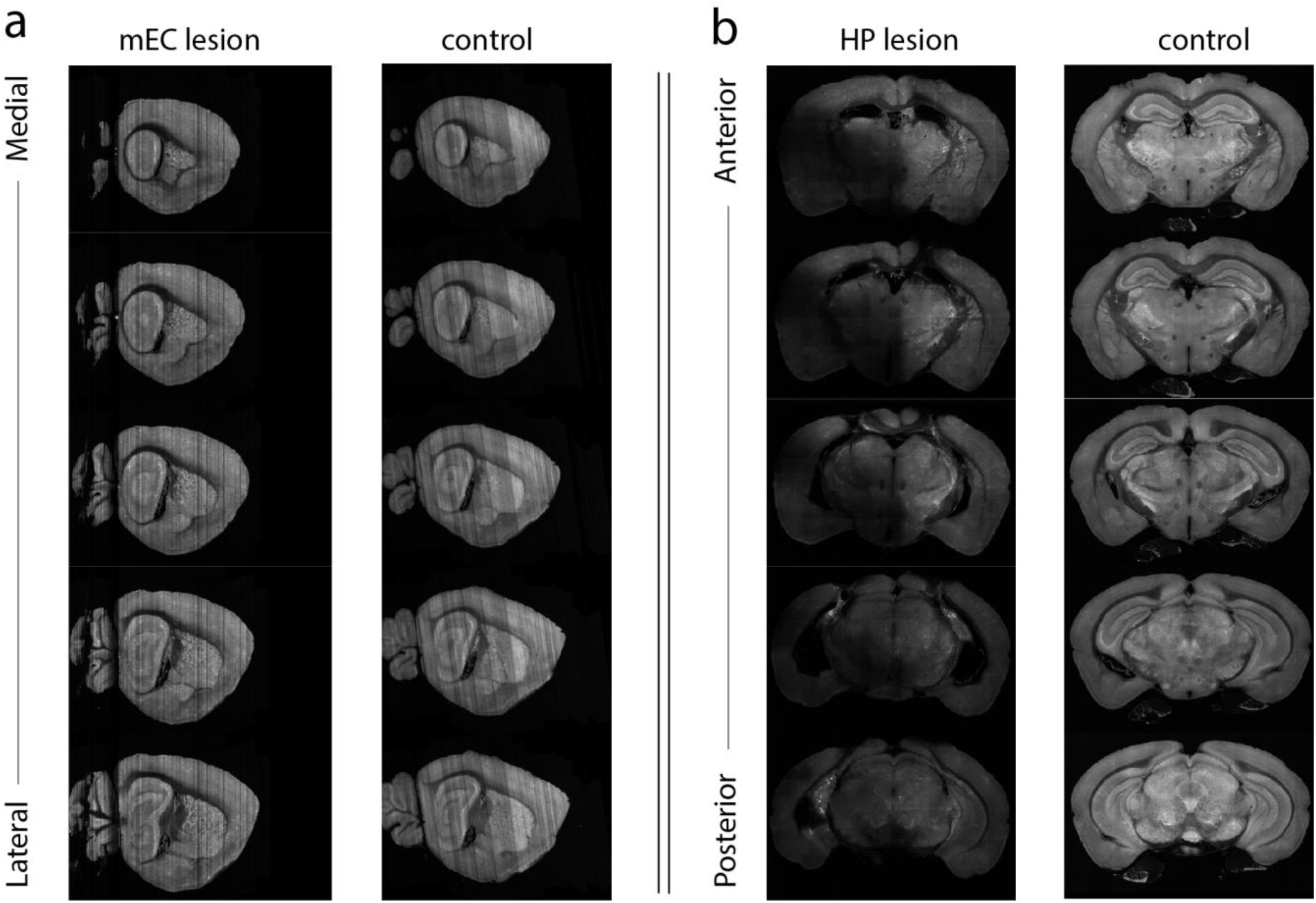
Histology. **(a)** An example mEC lesion. **(b)** An example HP lesion.

**Extended Data Fig. 3:**
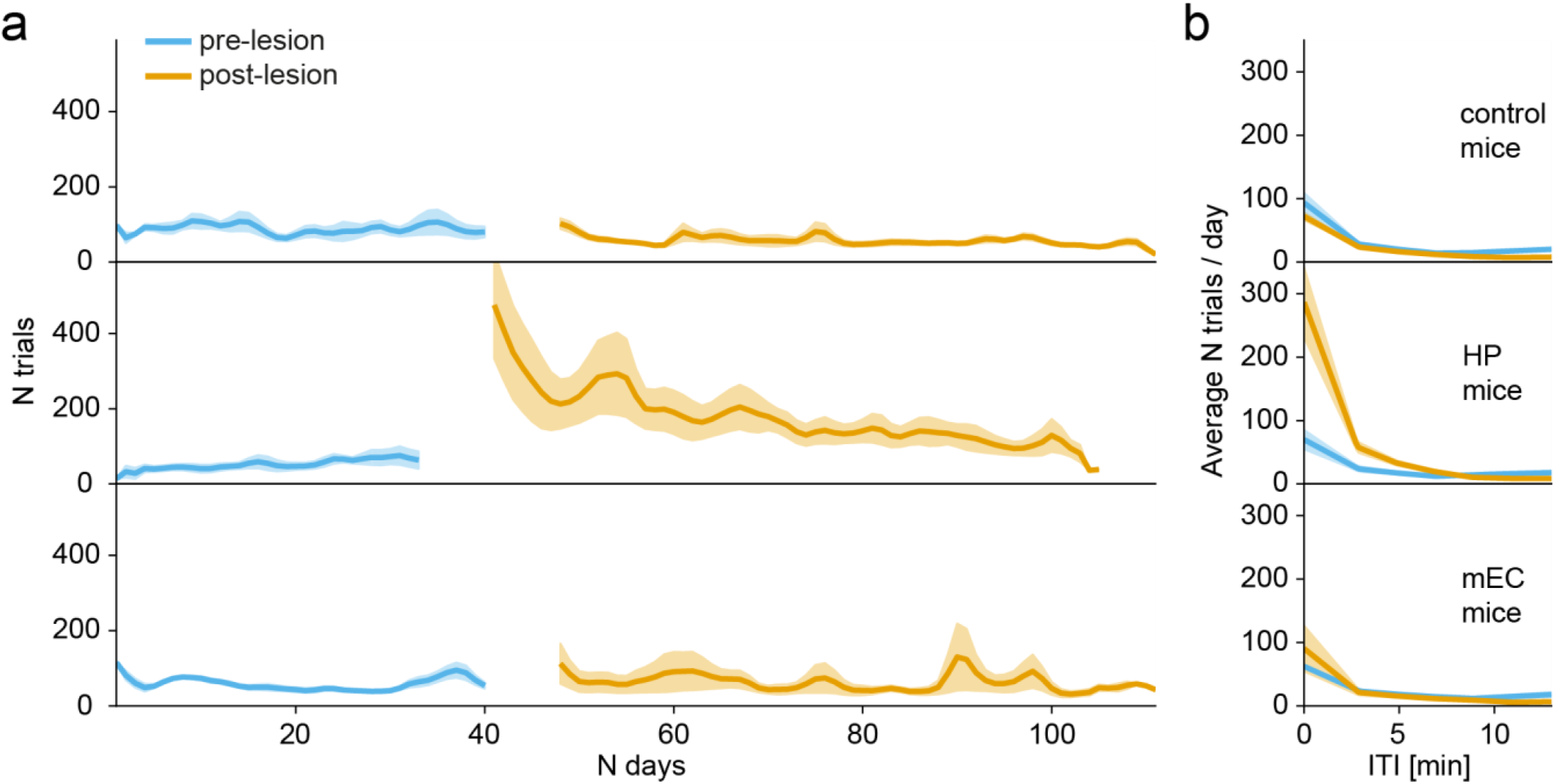
Nose-poking behaviour. **(a)** Average daily frequency of nose poking after an initial drinking attempt pre- (blue) and post- (orange) surgery in sham control (top), HP (middle) and mEC (bottom) mice. **(b)** Average daily frequency of noise-poking at different ITIs pre- (blue) and post- (orange) surgery in sham control (top), HP (middle) and mEC (bottom) mice.

**Extended Data Fig. 4:**
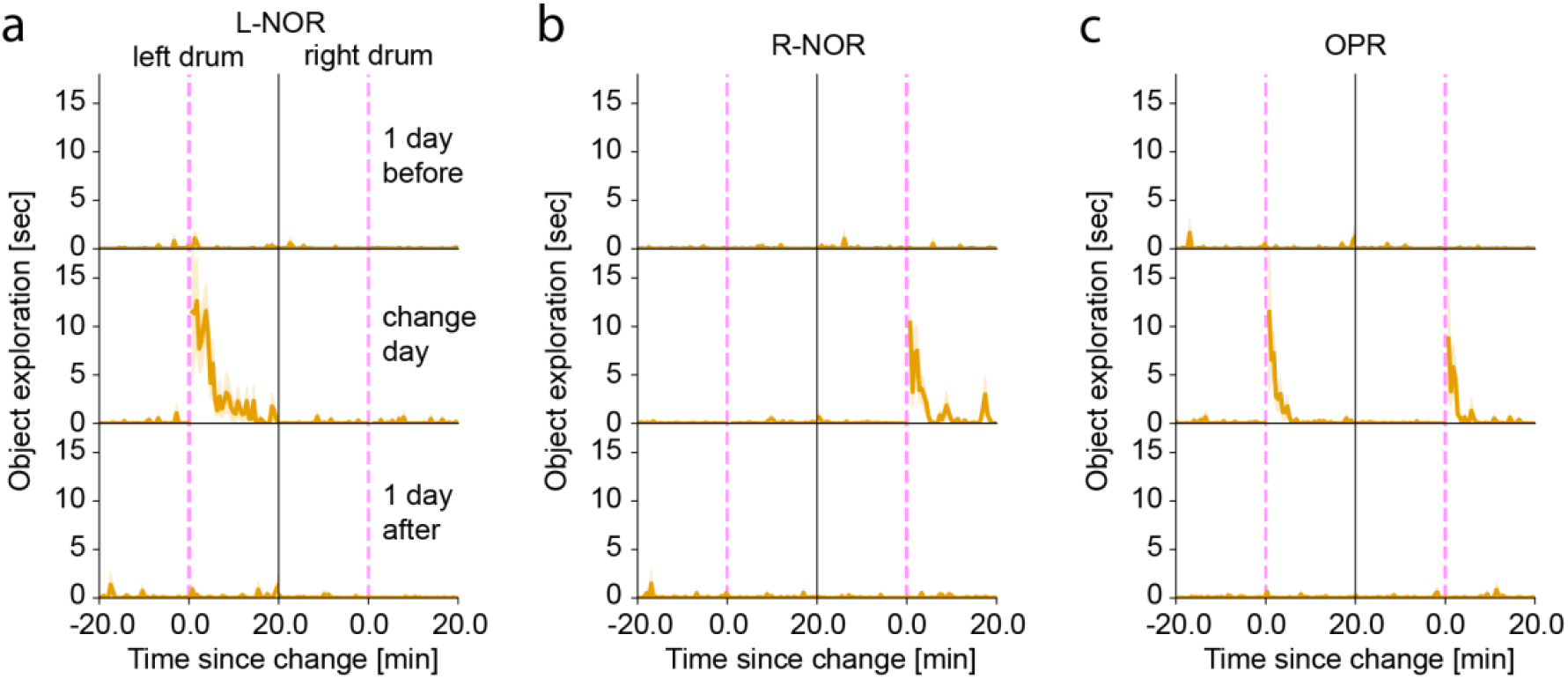
One-trial object and object-in-place recognition in the smart-Kage. **(a)** left-NOR task. **(b)** right-NOR task. **(c)** OPR task in pre-lesioned WT mice. Object exploration one day before the change (top row) measured at ∼12 pm, corresponding to the usual time of a side panel change on the day of the change (middle row) and one day after the change (bottom row). A dashed pink line indicates the time of the change.

**Extended Data Fig. 5:**
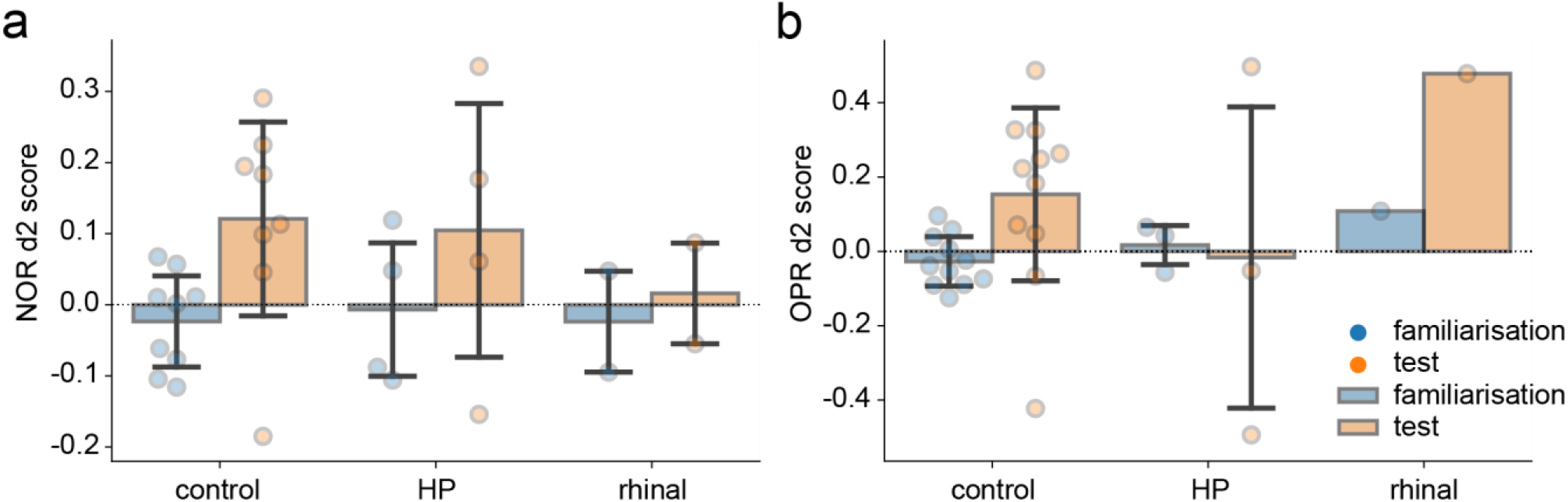
Standard novel object recognition and object-in-place recognition tests. **(a)** Standard NOR test. The differences between d2 scores during the test sessions are not significant between the groups (control vs. HP: U=17.0, P=0.9333; control vs. mEC: U=13.0, P=0.8000; HP vs. mEC: U=5.0, P=0.9333; Mann-Whitney U-test). **(b)** Standard OPR test. Similarly, the differences between d2 scores during the test sessions are not significant between the groups (control vs. HP: U=20.0, P=0.9890; control vs. mEC: U=1.0, P=0.9890; HP vs. mEC: U=1.0, P=1.0; Mann-Whitney U-test). Blue: familiarization session; orange: test session; dots - individual animals.

**Extended Data Fig. 6:**
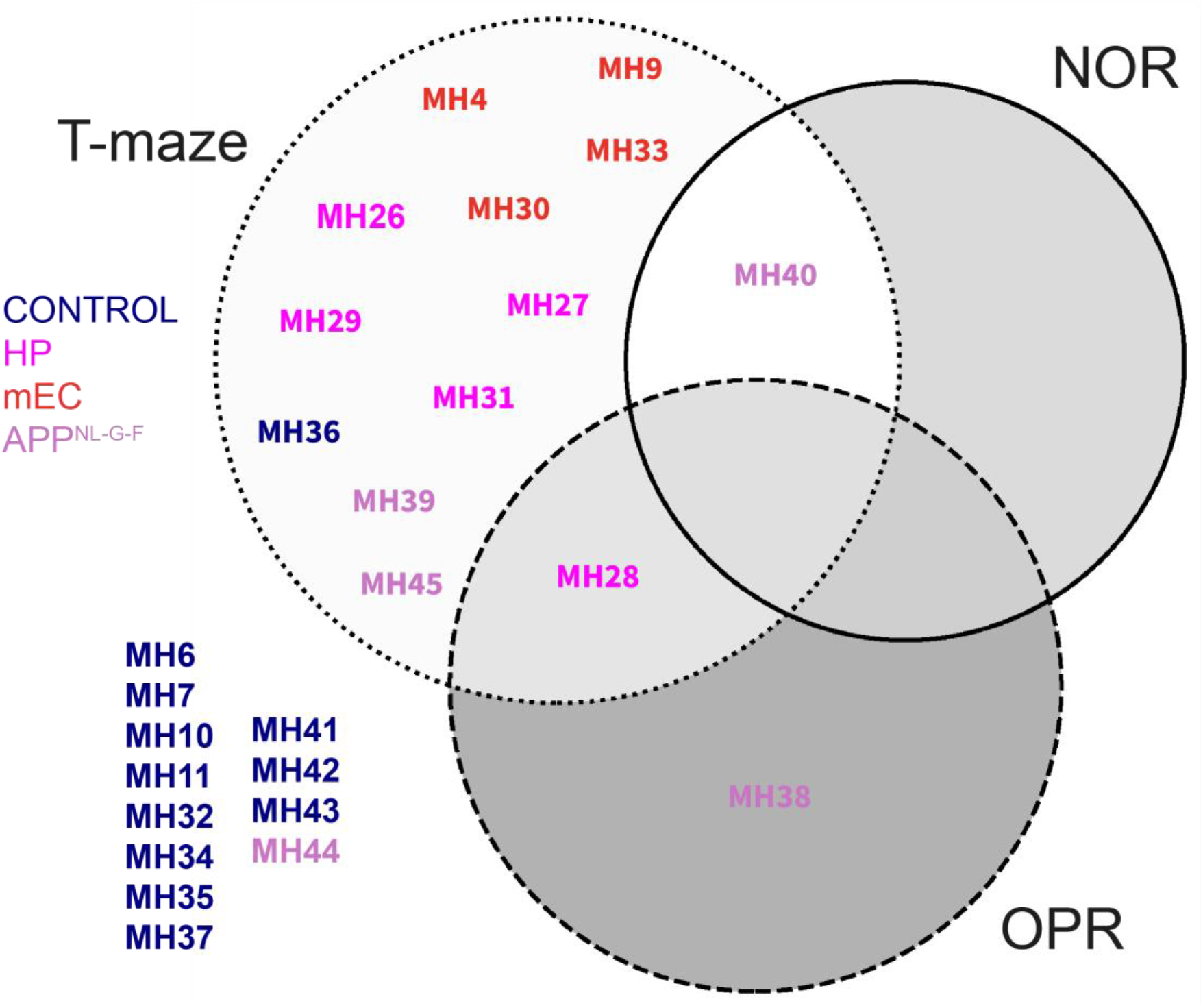
Mouse group identification on an individual animal basis from standard T-maze, novel object recognition and object-in-place recognition tests. Different colours represent a mouse group. The pie charts indicate mice that did not pass the high-performance threshold. The thresholds were selected to optimise the number of mice correctly identified as the control group shown outside the pie charts.

### Supplementary Information

Supplementary Fig. 1

Supplementary Tables 1-4

**Supplementary Video 1. An example of a mouse performance on the smart T-maze task overlaid with CNN tracking**.

**Supplementary Video 2. An example of a mouse exploring one of the side panels during a smart NOR task overlaid with CNN tracking**.

**Supplementary Video 3. An example of a mouse running on the wheel overlaid with CNN tracking**.

**Supplementary Video 4. An example of a mouse in a quiescence state overlaid with CNN tracking**.

**Supplementary Video 5. An example of a mouse with a hippocampal lesion running in a circular trajectory**. This behaviour was not observed in sham controls and mice with entorhinal lesions.

