## Supplementary Information for "A novel fully-automated system for lifelong continuous phenotyping of mouse cognition and behaviour"

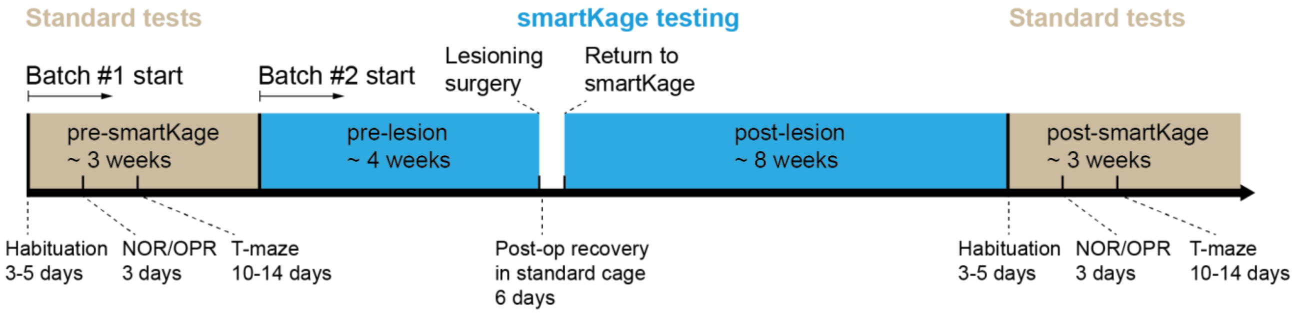


**Supplementary Figure 1: Experimental timeline for batch#1 & #2 smart-Kage experiment.**

**Supplementary Table 1. A basic drum sequence**. For longer testing periods, the sequence was approximately restarted.

| **Week** | **Day** | **Drum change** |
|---|---|---|
| 1 | 1 | both NOR |
| | 3 | left NOR |
| | 5 | left OPR, right NOR |
| | 7 | right NOR |
| 2 | 2 | left NOR, right OPR |
| | 4 | left NOR |
| | 6 | both OPR |
| 3 | 1 | both NOR |
| | 3 | left NOR, right OPR |
| | 5 | both NOR |
| | 7 | right NOR |
| 4 | 2 | left OPR, right NOR |
| | 4 | left NOR |
| | 6 | *repeat* |

**Supplementary Table 2. Identities of hippocampal- and medial entorhinal-lesioned animals and corresponding quantification of lesioned size.**

| Lesion site | Mouse ID | Average combined lesion size (%) | Left hemisphere lesion size (%) | Right hemisphere lesion size (%) |
|---|---|---|---|---|
| HP | MH26 | 56 | 67 | 44 |
| | MH27 | 52 | 58 | 46 |
| | MH28 | 52 | 38 | 66 |
| | MH29 | 67 | 82 | 52 |
| | MH31 | 49 | 54 | 43 |
| mEC | MH04 | 20 | 12 | 28 |
| | MH09 | 65 | 92 | 38 |
| | MH30 | 97 | 97 | 97 |
| | MH33 | 52 | 60 | 44 |

**Supplementary Table 3. Mouse groups tested in the smart-Kage.**

| Group name | Lesion or genotype | Number of mice | Age at the start in the smart-Kage (weeks) | Gender |
|---|---|---|---|---|
| Batch #1 | Sham | 4 | 10 | M |
| | mEC | 2 | 10 | M |
| Batch #2 | Sham | 5 | 16 | M |
| | mEC | 2 | 16 | M |
| | HP | 5 | 16 | M |
| *APP* | control | 3 | 22 | M |
| | mutant | 5 | 22-24 & 39 | M |

**Supplementary Table 4. Behavioural features used in unsupervised clustering.**

| Category | Features |
|---|---|
| T-maze | maximal daily-average trial frequency w.r.t ITI |
| | daily-average trial frequency mean |
| | daily-average trial frequency slope [1/day] |
| | maximal T-maze performance w.r.t ITI [%] |
| | ITI at middle point between maximal and 50% T-maze performance [min] |
| | ITI at 50% T-maze performance [min] |
| NOR | average maximum exploration after drum change [min] |
| | average elapsed time between drum change and maximal exploration [min] |
| | average elapsed time between drum change and half-maximal exploration [min] |
| OPR | average maximum exploration after drum change [min] |
| | average elapsed time between drum change and maximal exploration [min] |
| | average elapsed time between drum change and half-maximal exploration [min] |
| Wheel Running | daily-average time of running on the wheel mean [min] |
| | daily-average time of running on the wheel slope [min/day] |
| Quiescence | <u>Light phase:</u> |
| | daily-average quiescence frequency mean |

daily-average quiescence frequency slope [1/day]

daily-average quiescence time mean [min]

daily-average quiescence time slope [min/day]

Dark phase:

daily-average quiescence frequency mean

daily-average quiescence frequency slope [1/day]

daily-average quiescence time mean [min]

daily-average quiescence time slope [min/day]

Combined:

daily-average total quiescence time mean [min]

daily-average total quiescence time slope [min/day]
